# High-throughput Single-Cell Proteomics and Transcriptomics from the Same Cells with a Nanoliter-Scale Spin-Transfer Approach

**DOI:** 10.1101/2025.10.20.681067

**Authors:** Pranav Dawar, Lye Meng Markillie, Sarah M. Williams, Hugh D. Mitchell, Johannes W. Bagnoli, Joshua Cantlon-Bruce, Anjali Seth, Carter C. Bracken, Ljiljana Paša-Tolić, Ying Zhu, James M. Fulcher

## Abstract

Single-cell multiomic platforms provide a comprehensive snapshot of cellular states and cell types by offering critical insights into the spatiotemporal regulation of biomolecular networks at a systems level, thereby defining the basis of multicellularity. Here, we introduce nanoSPINS, an advanced platform that enables high-throughput profiling and integrative analysis of the transcriptome and proteome from the same single cells using RNA sequencing and isobaric labeling LC-MS-based proteomics, respectively. NanoSPINS can efficiently transfer mRNA-containing droplets across two microarrays via a centrifugation-based approach, while proteins are retained on the initial platform. Benchmarking of nanoSPINS on two cell lines demonstrates its ability to generate global proteomic and transcriptomic profiles that align well with previously established methodologies/platforms. The incorporation of isobaric TMTpro labeling into this single-cell multiomics platform significantly enhances the throughput of single-cell proteomic analyses. Through the high-throughput quantification of the proteome and transcriptome, nanoSPINS not only facilitates the identification of molecular features at both mRNA and protein level but also provides larger sample sizes for improved statistical power in clustering and differential abundance. Given the broad applicability of single-cell multiomics in biological research and clinical settings, we believe nanoSPINS represents a powerful platform for the characterization of heterogeneous cell populations.

## INTRODUCTION

Multicellular systems exhibit significant complexity, encompassing a variety of cell types with distinct morphological, anatomical, and functional characteristics. This complexity manifests at both intercellular (between different cell types) and intracellular (between different biomolecules) levels. As the differential regulation of biomolecules is fundamental to multicellularity, the quantification of these biomolecules (particularly mRNAs and proteins) is crucial for describing cell types, cell states, and overall understanding of cellular responses to various stimuli. The rapid advancement of single-cell omic technologies, including both sequencing-based and mass spectrometry-based methods, have significantly enhanced our ability to capture these cellular facets. Over the past decade, “global” single-cell transcriptomics (scRNAseq) methods have been optimized to provide remarkable improvements in sensitivity, throughput, and cost-effectiveness^1, 2^. Mass-spectrometry-based global single-cell proteomics (scProteomics) has seen similar advances while also facing unique challenges, such as sensitivity and throughput^3^. To address these challenges, substantial efforts have been put forth in miniaturizing sample preparation steps, optimizing analytical workflows, and applying new mass spectrometers^4–6^. Despite these advances, singular measurements of individual classes of biomolecules within a cellular system only provide a partial view of the cell; hence multiomic measurements to connect modalities are paramount to a holistic understanding.

Several recent developments have enabled simultaneous or parallel multimodal omics profiling at single-cell resolution. However, most of these approaches are limited to sequencing-based platforms that co-profile the transcriptome and epigenome, such as scMT-seq^7^, snmCT-seq^8^, scTrios-seq^9^, and scNOMeRe-seq^10^. Antibody-based sequencing platforms have also been developed to measure a small set of proteins and the whole transcriptome, such as CITE-seq^11, 12^ and DBiT-seq^13^. The recent introduction of nanoSPLITS (nanodroplet SPlitting for Linked-multimodal Investigations of Trace Samples) enables the same single-cell, parallel measurement of the transcriptome and the proteome using scRNAseq and mass-spectrometry-based proteomics, respectively^14^. This platform achieves equal splitting of nanoliter-scale droplets containing cellular material, followed by low-volume proteomic and transcriptomic sample preparation. By miniaturizing assay volumes to nanoliter scales, sample losses due to non-specific adsorption are minimized and tryptic digestion kinetics are enhanced, leading to improvements in identification depth. While feasibility is established and highly quantitative accuracy is obtained, one practical disadvantage of nanoSPLITS approach is relatively low throughput due to the use of label-free proteomics workflow, particularly when compared to scRNAseq approaches, which can routinely measure tens to hundreds of thousands of cells in a single sequencing experiment.

Notably, several multiplexing scProteomic approaches based on non-isobaric mass tags (e.g., plexDIA^15, 16^, mTRAQ^16^) and isobaric mass tags (e.g., TMT/TMTpro^16–19^) have been developed. The integration of these multiplexing approaches has the potential to increase the overall throughput of single cell multiomics by 3-32 times, depending on plexing capability and other factors. Our group has developed a high-throughput and nanoliter-scale multiplexed scProteomics technology, referred to as nested nanoPOTS or N2^18^, for scalable measurements of single cells. Importantly, the nested well design significantly improved the nanowell density, enabling the integration of more than 500 wells in a single glass chip.

Herein, we describe an improved sample preparation workflow that enables high-throughput same-cell scProteomics and scRNA-seq by integrating TMT-based multiplexed proteomics with barcode-based scRNA-seq. Instead of relying on manual droplet splitting in our nanoSPLITS approach, we developed a spin-transfer approach (which we refer to as “nanoSPINS”), that utilizes centrifugation to separate the mRNA and proteins into two different devices. NanoSPINS can efficiently retain protein on custom-designed nanoSPINS chip while transferring mRNA-containing droplet into the cellenCHIP 384 for barcoded scRNAseq analysis. We demonstrate the nanoSPINS-based multiomics platform can offer comprehensive molecular characterizations by quantifying cell-type-specific features across modalities and expanding applications beyond the reach of single-omics measurements alone.

## MATERIALS AND METHODS

### Reagents and chemicals

Deionized water (18.2 MΩ) was purified using a Barnstead Nanopure Infinity system (Los Angeles, CA, USA). n-dodecyl-β-D-maltoside (DDM), iodoacetamide (IAA), ammonium bicarbonate (ABC), and formic acid (FA) were obtained from Sigma (St. Louis, MO, USA). Nuclease-free water (not DEPC-treated), mass spectrometry grade Trypsin Platinum (Promega, Madison, WI, USA) and Lys-C (Wako, Japan) were reconstituted in 50 mM ABC before usage. Dithiothreitol (DTT, No-Weigh format), acetonitrile (ACN) with 0.1% FA, water with 0.1% FA (MS grade), anhydrous dimethylsulfoxide, Hoescht 33342 stain, and ATTO-TAG FQ kit were purchased from Thermo Fisher Scientific (Waltham, MA, USA). cellenCHIP 384-3’ RNAseq kit was acquired from Cellenion.

### NanoSPINS and nanoSPLITS chip design and fabrication

NanoSPLITS and nanoSPINS chips were fabricated using standard photolithography, wet etching, and silanization as described previously^5^. Chip designs were produced using AutoDesk AutoCAD 2022. The nanoSPLITS chips contained 48 (4 ×12) nanowells with a well diameter of 1.2 mm and inter-well distance of 4.5 mm. The nanoSPINS chips contained 4 arrays of 96 wells (384 total) in a 12 × 8 format with a well diameter of 0.6 mm and interwell distance 1.5 mm. Chip fabrication utilized a 25 mm x 75 mm glass slide pre-coated with chromium and photoresist (Telic Company, Valencia, USA). After photoresist exposure, development, and chromium etching (Transene), select areas of the chip were protected using Kapton tape before etching to a depth of ∼5 µm with buffered hydrofluoric acid. The freshly etched slide was dried by heating it at 120 °C for 1 h and then treated with oxygen plasma for 3 min (AP-300, Nordson March, Concord, USA). 2% (v/v) heptadecafluoro-1,1,2,2-tetrahydrodecyl-dimethylchlorosilane (PFDS, Gelest, Germany) in 2,2,4-trimethylpentane was applied onto the chip surface and incubated for 30 min to allow for silanization. The remaining chromium covering the wells was removed with etchant, leaving elevated hydrophilic nanowells surrounded by a hydrophobic background. For all chips, PEEK chip covers were machined to fit the chip and used to prevent droplet evaporation during sample handling steps. Chips were wrapped in parafilm and aluminum foil for long-term storage as well as intermediate steps during sample preparation.

### Cell culture and cellenONE sorting

Two murine cell lines (NAL1A clone C1C10 is referred to as C10 and is a non-transformed alveolar type II epithelial cell line derived from normal BALB/c mouse lungs; SVEC4-10, an endothelial cell line derived from axillary lymph node vessels) were cultured at 37°C and 5% CO2 in Dulbecco’s Modified Eagle’s Medium supplemented with 10% fetal bovine serum and 1× penicillin-streptomycin (Sigma, St. Louis, MO, USA). Cultured cell lines were collected in a 15 ml tube and centrifuged at 1,000 × g for 3 min to remove medium when preparing for cellenONE sorting. Cell pellets were washed three times by PBS, then counted to obtain cell concentration. PBS was then added to achieve a concentration of 2 × 10^6^ cells/mL. Immediately before cell sorting, the cell-containing PBS solution was passed through a 40 µm cell strainer (Falcon™ Round-Bottom Polystyrene Test Tubes with Cell Strainer Snap Cap, FisherScientific) to remove aggregated cells.

A cellenONE instrument equipped with a glass piezo dispensing capillary (P-20-CM) for dispensing and aspiration was utilized for single-cell isolation. Sorting parameters included a pulse length of 50 µs, a nozzle voltage of 80 V, a frequency of 500 Hz, a LED delay of 200 µs, and a LED pulse of 3 µs. The slide stage was temperature-controlled utilizing dew-point control mode to reduce droplet evaporation on nanoSPLITS and nanoSPINS chips. Cells were isolated based on their size, circularity, and elongation in order to exclude apoptotic cells, doublets, or cell debris. For C10 cells, this corresponded to 25 to 40 µm in diameter, maximum circularity of 1.15, and maximum elongation of 2, while SVEC cells were 24 to 32 µm in diameter, maximum circularity of 1.15, and maximum elongation of 2. For both cell types, and specifically in the nanoSPINS experiments, fluorescent staining was performed prior to sorting by incubating the samples with Hoescht 33342 (1:10,000 dilution) for 10 minutes to enable post-spin cell counting. All cells were sorted based on bright field and/or fluorescent images (where applicable) in real time.

### Fluorescent protein labeling experiments

For ATTO TAG FQ fluorescent experiments with nanoSPLITS chips, nanowells were first loaded with 200 nL 10 mM Tris (with or without 0.1% DDM). C10 cells (∼1 × 10^6^) were incubated in 0.9 mL of 1 mM 3-2-(furoyl quinoline-2-carboxaldehyde (FQ), 10% dimethylsulfoxide and 9 mM potassium cyanide with 45 mM triethylammonium bicarbonate at room temperature for 1 h to achieve labeling of primary amines on cellular proteins. Cells were pelleted at 1,000 × g for 5 minutes before reaction buffer was removed, followed by resuspension in 1 mL PBS and filtering through a 40 μm Corning cell strainer. Finally, this filtered cell solution was used for sorting on the cellenONE. Fluorescent imaging was performed on a PALM MicroBeam system (Carl Zeiss MicroImaging, Munich, Germany). Fluorescent images were generally collected at 10 to 20× magnification through a Zeiss FL 38 HE filter set for excitation of 450-490 nm and emission at 500-550 nm.

### NanoSPINS workflow to separate protein and mRNA from same cells

First, cellenCHIP 384 nanowell plates were loaded with 1× RT mix (100 nL for control wells) (Cellenion, cat# CEC-5015-8) or 1.7× RT mix (60 nL for wells that will receive lysate from single-cells) before being frozen at -80 °C until use. NanoSPINS chips were loaded with 40 nL of 10 mM HEPES pH 8.5 with 0.1% DDM, prepared either freshly or from frozen stock stored at –80 °C. Next, C10 and SVEC cells labeled with Hoescht 33342 stain were sorted using the cellenONE onto nanoSPINS chip wells, as well as positive control wells on the cellenCHIP 384. Furthermore, to create a TMT boost channel for the nanoSPINS samples, 30 cells (10 C10 and 20 SVEC) were sorted and pooled onto a separate standard nanoSPLITS chip. The nanoSPINS chips were then either directly centrifuged (1000 × g for 0.5 min) into the cellenCHIP 384 plates or allowed to evaporate at room temperature before reconstitution with 40 nL of nuclease-free water followed by centrifugation into the cellenCHIP. After centrifugation, nanoSPINS chips were imaged using a Zeiss Axio Zoom microscope (Carl Zeiss MicroImaging, Munich, Germany) with brightfield and a fluorescent DAPI filter set in order to quantify retention of cell nuclei on chips while cellenCHIP 384 plates were directly placed into the thermocycler for RT reaction. The nanoSPINS chips were stored at -80 °C before scProteomic sample preparation.

### scProteomic sample preparation and LC-MS/MS analysis

All nanoSPINS and nanoSPLITS chips were first allowed to dry before being placed onto the dewpoint-controlled deck of the cellenONE system for sample preparation. Protein extraction, reduction, and alkylation were accomplished in nanoSPINS chips via a one-step dispensing of 10 nL of 10 mM HEPES, 2 mM DTT, and 5 mM IAA before sealing the chip and incubating in a humidified box at 70°C for 30 min, followed by room temperature incubation for another 15 min. The nanoSPLITS chips containing carrier channel samples had 100 nL of 10 mM HEPES, 2 mM DTT, 5 mM IAA, and 0.1% DDM added before performing the same incubation steps as above. Digestion was accomplished through the addition of 5 nL water containing 0.25 ng trypsin and 0.075 ng Lys-C to nanoSPINS wells or 100 nL water containing 2.5 ng trypsin and 0.75 ng Lys-C for nanoSPLITS boost channel samples, followed by overnight digestion at 37°C. Prior to the addition of TMTpro reagents, the cellenONE deck was adjusted to a temperature of 18°C, and humidity was increased for evaporation control at a dewpoint near this temperature. TMTpro reagents were dissolved in anhydrous DMSO at a concentration of 5 ng/nL before the addition of 10 nL (50 ng per channel) to each nanoSPINS sample or 100 nL (500 ng) to carrier channel samples. Labeling was allowed to proceed for 1 hr at 18°C before addition of 2 nL 5% hydroxylamine (20 nL for carrier samples) and a further incubation of 15 min. A final quenching was accomplished through the addition of 5 nL 5% formic acid (50 nL for carrier samples) prior to drying the samples in a vacuum desiccator. All chips were then stored at -20 °C until LC-MS analysis.

For LC-MS analysis, we employed an in-house assembled nanoPOTS autosampler for LC-MS analysis ^20^. The autosampler contains a custom packed SPE column (100 μm i.d., 4 cm, 5 μm particle size, 300 Å pore size C18 material, Phenomenex) and an analytical LC column (50 μm i.d., 25 cm long, 1.7 μm particle size, 190 Å pore size C18 material, Waters) with a self-pack picofrit (cat. no. PF360-50-10-N-5, New Objective, Littleton, MA). The analytical column was heated to 50 °C using AgileSleeve column heater (Analytical Sales and services, Inc., Flanders, NJ). To collect samples from the nanoSPINS chips, the collection capillary was placed on the center well for each TMT plex before addition of 6 μL of Buffer A (0.1% formic acid in water). For each plex, three 6 μL collections (two for washing purposes) were performed before a boost channel well was collected using 400 nL of Buffer A. The entire volume was collected into a 25 μL sample loop before injection and trapping on the SPE column for 5 min. After washing the peptides, samples were eluted at 100 nL/min and separated using a 120-min gradient from 8% to 45% Buffer B (0.1% formic acid in acetonitrile).

An Orbitrap Fusion Lumos Tribrid MS (ThermoFisher Scientific) without FAIMS, operated in data-dependent acquisition (DDA) mode, was used for all analyses. Source settings included a spray voltage of 2,200 V and ion transfer tube temperature of 200°C. Orbitrap MS1 acquisitions were performed across the 450-1600 m/z range with a maximum injection time of 254 ms and AGC target of 250% (1e6 charges) at 120,000 resolution. For DDA MS2 acquisitions, charge states 2-7 were considered, MIPS mode was set to “Peptide”, intensity threshold was set to 20,000, and dynamic exclusion was set to 50 s within a 10-ppm tolerance of the precursor. Orbitrap MS2 acquisitions were performed with 35% HCD at 120,000 resolution, a maximum injection time of 246 ms, and an AGC target of 2,000 % (1e6 charges).

### scRNAseq library preparation and Sequencing

Following the transfer of samples into RTready cellenCHIP 384 (Cellenion, cat# CEC-5015-8) with each well preloaded with RT buffer containing a uniquely barcoded oligo(dT) primer. The reverse transcription reaction was performed at 42 °C for 90 minutes to generate the first strand cDNA. Following the first strand cDNA synthesis, all samples from cellenCHIP 384 was pooled into one reaction tube for cDNA amplification to generate the second strand of cDNA. 1 ng of the double strand cDNA was fragmented and tagged with specific primer using cellenTAG reagent and cellenTAG enzymes (Cellenion, cat# CEC-5015-8). The tagged cDNA was subsequently amplified using 12 PCR cycles to enrich the template libraries suitable for sequencing on the illumina platform. The resulting libraries were sequenced on the illumina NextSeq 550 system using NextSeq 500/550 High Output v2 kit 150 cycles (cat#20024907) with a paired-end sequencing configuration consisting of 26 cycles for Read 1, 8 cycles for i7 Index, and 100 cycles for Read 2, according to cellenCHIP 384 - 3’RNA-seq Kit sequencing guidelines. Data quality was assessed with FastQC, and quality ensured raw fastq datasets were processed using zUMIs v2.9.3e ^21^ in combination with STAR v2.7.3a ^22^. Reads were mapped to the mouse (mm10) reference genome and gene annotations were obtained from gencode.vM25 (GRCm38.p6). Additionally, down sampling to fixed numbers of raw sequencing reads per cell was performed using the “-d” option in zUMIs.

### Data analysis

All raw files were processed by FragPipe^23, 24^ (version 22) and searched against the *Mus musculus* UniProt protein sequence database with decoy sequences (UP000000589 containing 17,196 forward entries, accessed 02/24). Search settings included a precursor mass tolerance of +/− 20 ppm, fragment mass tolerance of +/− 20 ppm, deisotoping allowing C12/C13 isotope errors of −1/0/1/2/3, strict trypsin and Lys-C as the enzyme (with allowance for N-terminal semi-tryptic peptides), carbamidomethylation and lysine TMT labeling (+304.20715) as a fixed modification, and several variable modifications, including oxidation of methionine, N-Terminal TMT labeling (+304.20715) and N-terminal acetylation. Protein and peptide identifications were filtered to a false discovery rate of less than 0.01 within FragPipe. TMT quantification was extracted from MS/MS spectra using philosopher-v5.1.1 and the output files were then processed using tmt-integrator-5.0.9 ^23^, to generate summary reports at the protein and peptide levels. FragPipe result files were then imported into RStudio for downstream analysis in the R environment (version 4.3.1). Labelling annotation of C10 and SVEC cells can be found in **Supplementary File 1**. Median normalization was performed with the “median_normalization” function from the proDA R ^25^ package across all scProteomics datasets. With regards to quality control (QC) filtering of scProteomics and scRNAseq data, cells with total identifications of less than 750 and 1500, respectively, were removed from analysis.

Additionally, all the scRNAseq datasets with less than 50 percent reads mapped to the reference transcriptome were filtered out. The ComBat algorithm with generalization for missing values was applied to correct TMT batch effects. In cases where complete data was needed, imputation for scProteomics data was performed with k-nearest neighbors’ imputation with < 50% missing values. For scRNAseq datasets, additional filtering criteria of a minimum of 1 raw read in at least 5 distinct cells for each cell-type was used to filter genes with stochastic abundance across C10 and SVEC cells. Additionally, the raw counts were normalized using calcNormFactors and cpm functions from edgeR package ^26^. Quality-controlled datasets were then used for principal component analysis (PCA) using PCAtools R package ^27^ and hierarchical clustering using corrplot R package. Gene Ontology enrichment analysis was performed with the gprofiler2 R package ^28^. Integration of scRNA-seq and scProteomics datasets, as well as identification of cell-type markers was performed Seurat R package ^29, 30^. When relevant, all raw p-values determined through statistical testing were corrected for multiple hypothesis testing using the Benjamini-Hochberg method.

## RESULTS

### Principles and design of the nanoSPINS approach

To increase the throughput of single cell Multiomics (scMultiomics), nanoSPINS builds and integrates two technologies, including nested nanoPOTS^18^ for multiplexing scProteomics and cellenCHIP 384^31^ for multiplexing scTranscriptomics analysis. The overall workflow is illustrated in **Figure 1**. Briefly, single cells were first sorted on nanoSPINS chip preloaded with cell lysis and permeabilization buffers using CellenONE. The cells were permeabilized to release mRNA inside the droplets, while most of the protein content was retained inside the cell pellet. The droplets were allowed to evaporate by adjusting the chip temperature, which were then rehydrated with RNase-free water. Next, the nanoSPINS chip containing single-cell lysate was aligned with cellenCHIP 384, followed by high-speed centrifugation to spin down the mRNA-containing droplets from nanoSPINS chip wells to cellenCHIP 384 wells. Finally, single-cell samples in the cellenCHIP 384 wells were processed using the cellenCHIP 384-3’ RNA-Seq Kit^31^ to generate high-throughput multiplexed scTranscriptomics data, while the single-cell pellets retained on nanoSPINS chips were subjected to tryptic digestion and TMTpro labeling to generate high-throughput multiplexed scProteomics data.

**Figure 1.**
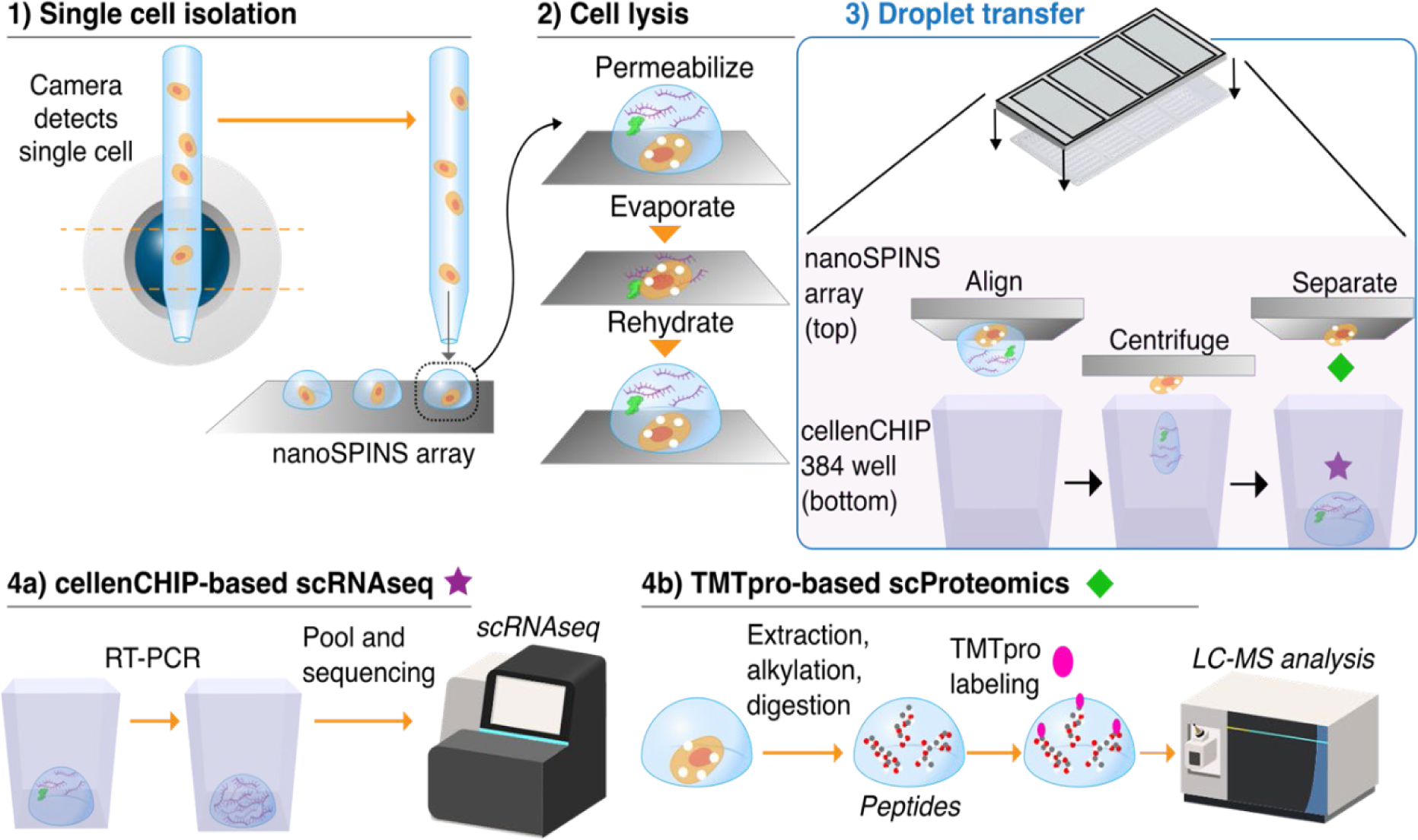
Schematic illustration of the nanoSPINS workflow, including cell sorting, permeabilization, droplet evaporation, rehydration, and spin-transfer of the cellular material for downstream scTranscriptomic with RNAseq and scProteomics with LC-MS.

### Large-scale spin-transfer of nanoliter droplet array from nanoSPINS chip to cellenCHIP-384

In our prior scMultiomics work with nanoSPLITS^14^, mRNA and protein contents are separated into two arrays by merging the two sessile droplets together, followed by manual splitting. While the nanoSPLITS workflow is simple, it is not compatible with the cellenCHIP format, where the nanowells are recessed. We therefore wondered if droplets could be transferred via centrifugation between the planar nanoSPINS chip and the cellenCHIP 384. Towards this end, we designed a prototype nanoSPINS chip with well-to-well dimensions matching those of the cellenCHIP (**Figure 2A**) while also incorporating a 3×3 arrangement for isobaric labeling of up to 9 cells with TMTpro (**Figure 2B**).

**Figure 2.**
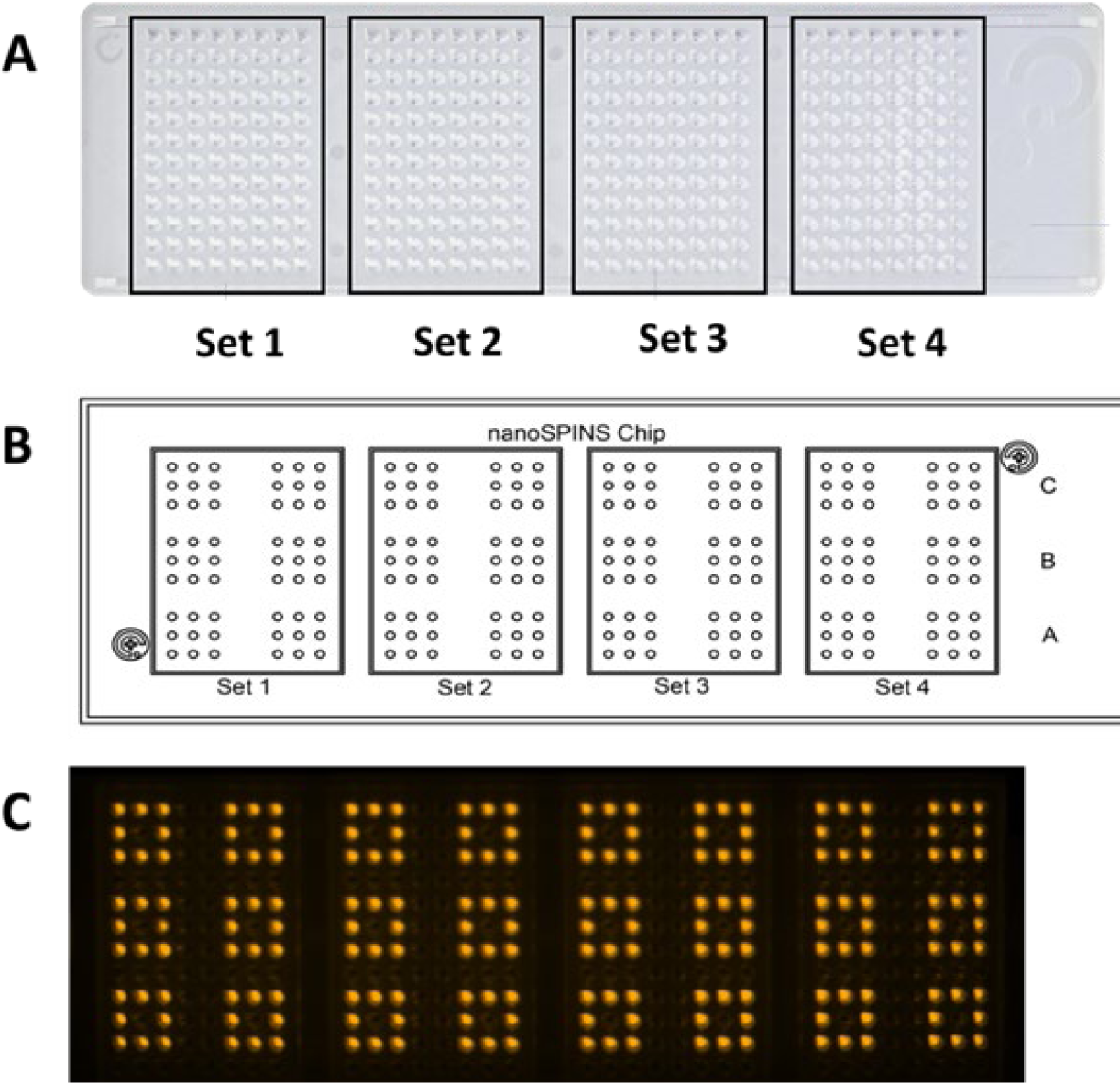
**(A)** Design of the cellenCHIP 384 showing the four sets of 96 wells. **(B)** nanoSPINS chip design with well dimensions matched to that of the cellenCHIP 384 and 3×3 arrangement for isobaric labeling of up to 9 cells in one batch. **(C)** Fluorescent image of the cellenCHIP 384 demonstrating successful centrifugation-based transfer (1000 × g) of 40 nL fluorescein-containing cell extraction buffer (0.000125% fluorescein, 10 mM Tris, 0.1% DDM) from the outer wells i.e., all eight peripheral wells of each 3×3 array (24 arrays in total) on the nanoSPINS chip. The central well in each 3×3 array, containing 40 nL of extraction buffer without fluorescein, served as a negative control.

Similar to previous iterations of nanoPOTS and nanoSPLITS chips, the wells are hydrophilic (having a bare silanol surface), while the areas surrounding the wells are modified to provide a hydrophobic boundary. We dispensed 40 nL of fluorescein-containing cell extraction buffer (0.000125% fluorescein, 10 mM Tris, 0.1% DDM) onto the outer 8 wells of the nanoSPINS chip, while the middle well had 40 nL of extraction buffer without fluorescein as a negative control. The droplet array was then manually aligned and adhered face-down to the cellenCHIP 384 with a standard PCR plate seal and was subjected to centrifugation at 1,000 × g for 1 min. Encouragingly, all 192 wells could successfully be transferred into the cellenCHIP 384 without cross-contamination of the negative control droplet or adjacent wells (**Figure 2C**).

### DDM-mediated permeabilization facilitates mRNA release while retaining proteins in the cell pellet

With successful proof-of-principle results in hand, we next turned our attention to the more challenging problem of getting cellular material (mRNA and protein) to be present in both the solution phase (i.e., nanodroplet) and on the solid phase (i.e., chip surface). In the previous nanoSPLITS workflow, a combination of a hypotonic solution (10 mM Tris), 0.1% DDM, and a freeze-thaw cycle at -80 °C was selected to minimize the use of reagents that might potentially interfere with downstream scRNA-seq or scProteomics, while also ensuring sufficient cell lysis. We found that a larger proportion (∼75%) of the protein content was retained on the chip originally containing the sorted cells^14^. The cause of this effect was not known but speculated to be related to protein diffusion or hydrophobic interactions of protein molecules with the trimethylsilyl-coated surface of the nanoSPLITS chip (**Supplementary Figure 1A**).

To provide better insights into cell lysis and protein retention, we replicated our original nanoSPLITS workflow by sorting pools (n = 10 and n = 5) of ATTO-TAG FQ-labeled C10 cells onto a nanoSPLITS chip containing 200 nL of nanoSPLITS lysis buffer (10 mM Tris, pH 7.5, 0.1% DDM) prior to a single freeze-thaw cycle at -80 °C. On the same chip, 200 nL of nanoSPLITS lysis buffer without 0.1% DDM was also included to investigate the impact of detergent on cell lysis. ATTO-TAG FQ is an amine-reactive for protein quantification, which becomes highly fluorescent after it reacts with primary amine. Using fluorescent microscopy imaging, diffuse or cloudy fluorescence was observed surrounding the C10 cell bodies or nuclei when DDM detergent was not added (**Figure 3A**). In contrast, the addition of DDM did not result in the same diffuse fluorescence surrounding the cells (**Figure 3B**). This observation suggested to us that hypotonic solution-induced ruptures of cell membranes in the absence of detergent were larger than with detergent, as the detergent-induced permeabilization reduces intracellular osmotic stress^32^. It is also notable that, independent of detergent, the cells remain mostly intact on the surface of the glass chip, as demonstrated by the bright areas of fluorescence representing cell bodies. This imaging data therefore helps to explain our previous observation of ∼75% of the protein content being retained on the array that cells were originally lysed. Therefore, our revised model of cellular lysis suggests cells can be permeabilized with 0.1% DDM, allowing diffusible cytosolic components such as mRNA and protein to move into the droplet while the bulk of the cell remains intact on the surface of the glass chip (**Supplementary Figure 1B**). Using the same experimental design with ATTO-TAG FQ-labeled C10 cells on nanoSPLITS chips with 200 nL of extraction buffer (with and without 0.1% DDM), we next tested the impact of centrifugation on the retention of cell pellet on chip surface. Interestingly, the presence of 0.1% DDM significantly enhanced the retention of cellular material on the chip (representative images shown in **Figures 3C** and **3D**). Across 66 cells tested without DDM, only 30 (45%) remained after centrifugation. In contrast, the presence of 0.1% DDM led to the retention of 64 out of 69 cells (93%). Although this data suggests DDM could be used to help retain cells on silanol surfaces, we also noted that the presence of DDM increased the droplet evaporation rate. Considering DDM also aids in reducing protein loss from non-specific adsorption in scProteomic experiments^33^, we next tested the effect of evaporation on the protein retention using 0.1% DDM as the cell lysis and permeabilization buffer.

**Figure 3.**
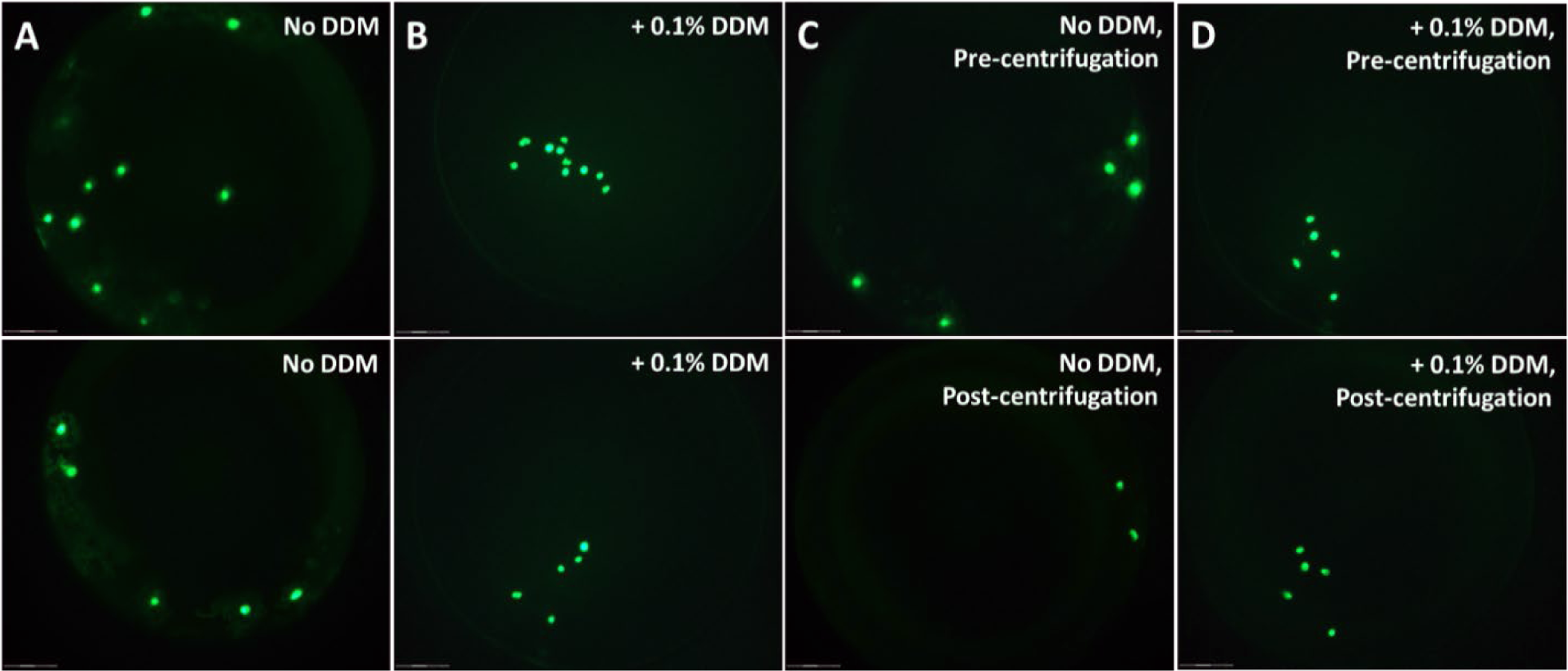
**(A)** Fluorescent image of pools of 10 (top) and 5 (bottom) ATTO-TAG FQ-labeled C10 cells sorted onto individual wells of a nanoSPLITS chip containing 200 nL of lysis buffer (10 mM Tris, without 0.1% DDM) exhibit diffuse or cloudy fluorescence around the cells, indicating low cellular integrity. **(B)** Fluorescent image of pools of 10 (top) and 5 (bottom) labeled C10 cells sorted onto wells containing lysis buffer with 0.1% DDM display bright, well-defined fluorescence, reflecting improved membrane integrity and enabling diffusion-mediated release of cellular contents into nanodroplets. **(C)** Fluorescent images of a pool of 5 labeled C10 cells sorted into lysis buffer without 0.1% DDM, before and after centrifugation at 1000 × g for 1 minute, reveal poor retention of cells on the nanoSPLITS chip. **(D)** In comparison, a pool of 5 labeled C10 cells sorted into lysis buffer with 0.1% DDM shows successful cell retention on the chip following the same centrifugation protocol.

Towards this end, we sorted a set of 192 C10 and SVEC cells (96 of each, labeled with Hoescht stain) onto three nanoSPINS chips, each, with wells containing 40 nL 0.1% DDM 10 mM HEPES pH 8.5. With two of the chips, we proceeded directly to centrifugation of the cell lysate into a cellenCHIP 384 after sorting and a single freeze-thaw cycle at -80 °C. For the third nanoSPINS chip, we allowed the droplets to fully evaporate and then reconstituted the wells with 40 nL of nuclease-free water, followed by centrifugation into a cellenCHIP 384. Surprisingly, for the two chips without an evaporation step, only 18.5% of the 384 cells could be observed with intact nuclear stain on the nanoSPINS chip. When evaporation was introduced (the third chip), the number of cell nuclei retained increased to 99% (190/192 cells, representative wells shown in **Supplementary Figure 2**), demonstrating that evaporation plays a key role in retaining non-solubilized cellular material onto the glass surface of the nested nanoPOTS chip. Therefore, the optimized nanoSPINS approach incorporates several key steps to achieve asymmetric splitting of cellular material, including the use of a 0.1% DDM-based cell lysis buffer and the incorporation of droplet evaporation and reconstitution steps before droplet spin-down (**Figure 1**).

### Optimized nanoSPINS workflow enables parallel characterization of two cell populations

With the approach fully optimized, we proceeded with downstream TMT-based scProteomic analysis of the single cells retained on the nanoSPINS chip and scRNAseq analysis of the transferred cell lysate in the cellenCHIP-384 containing the transferred cell lysate. We also included directly-sorted C10 and SVEC cells in cellenCHIP as positive control and empty wells as negative control (**Supplementary Figure 3**). From each TMTpro 16 plex, nine channels were used, with cells alternating such that there were four channels with C10 cells, four channels with SVEC cells, and a single blank channel (full sample arrangement on cellenCHIP 384 shown in **Supplementary Figure 3).**

TMTpro-based scProteomic analysis identified a total of 2,161 unique proteins (**Supplementary File 2**). After filtering 32 cells with fewer than 750 protein identifications, we retained a total of 160 cells, with 69 C10 cells and 91 SVEC cells. The QC-passed cells yielded an average of 5,066 peptides and 1,186 protein identifications (**Supplementary Figure 4A).** A minimum threshold of 750 proteins (**Supplementary Figure 4B**) was established based on the number of protein IDs detected in the blank channels, which can be attributed to isotope contamination across TMT channels, where ions from highly abundant samples may bleed into the reporter ion regions of adjacent channels, including blanks, resulting in false identifications. An average of 1,185 proteins were identified from C10 cells, and 1,188 proteins from SVEC cells (**Figure 4A**). We observed high Pearson correlations (∼0.95) between the respective proteomes of C10 and SVEC single cells (**Supplementary Figure 4C**), indicating high reproducibility of the proteomic data generated using the nanoSPINS workflow. In agreement with this, median coefficients of variation (CVs) were 0.267 and 0.277 for C10 (n = 69) and SVEC (n = 91) cells (**Figure 4B**). Additionally, principal component analysis (PCA) indicated that the PC2 pseudo-dimension correlated significantly with cell type (pvalue < 0.001) and accounted for ∼9% of the observed variation, allowing for clear separation of cell types (**Figure 4C**).

**Figure 4.**
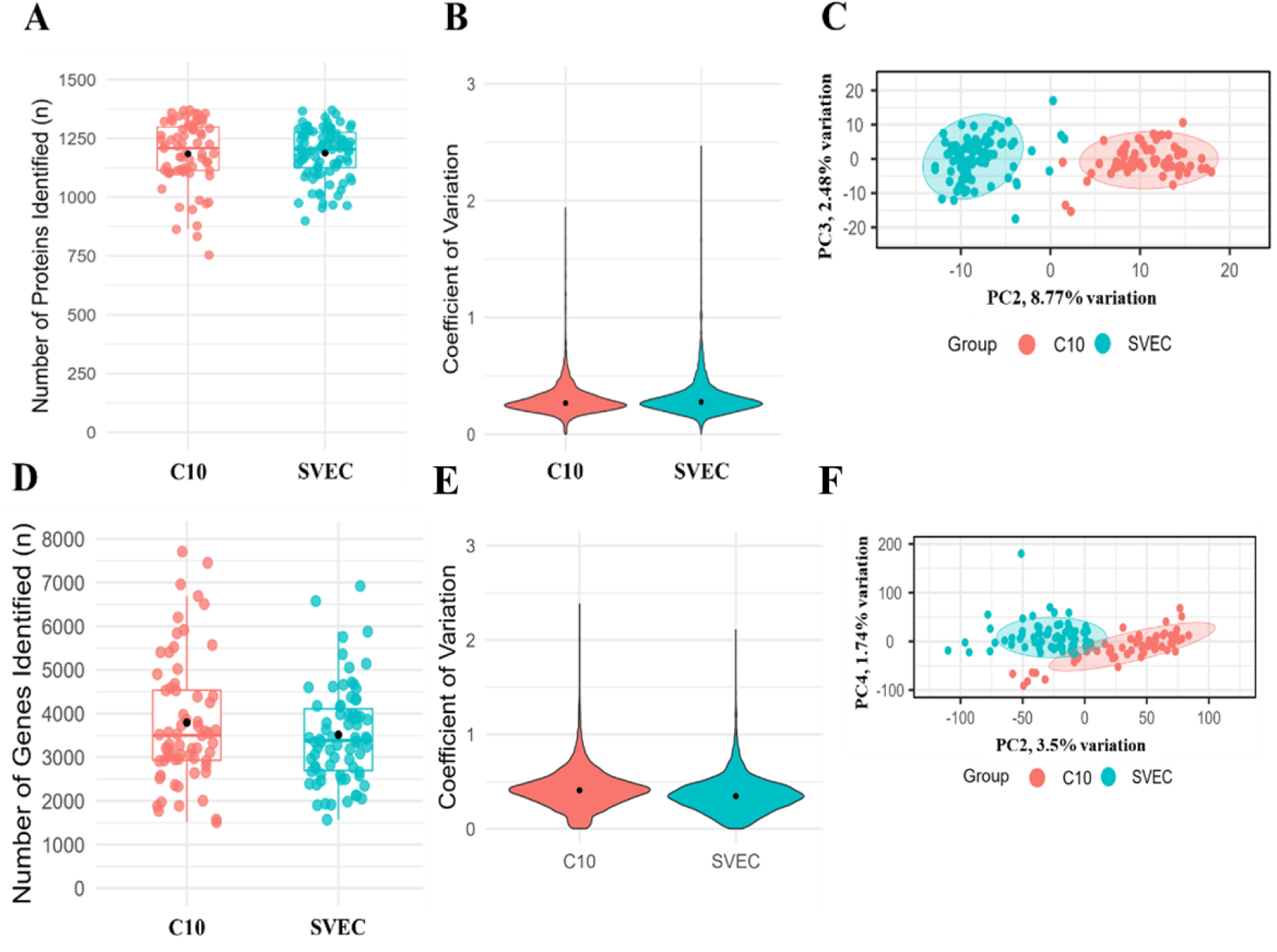
**(A)** Box plot showing the distributions of protein identification numbers for QC-passed 69 C10 cells and 91 SVEC cells. Centerlines represent the distribution median and “black” points represent the distribution mean. **(B)** Violin plots showing the coefficient of variations of protein abundances from QC-passed scProteomic samples processed using nanoSPINS (n = 2,098 proteins for C10 and n = 2,115 proteins for SVEC cells). “Black” point represents median CV. **(C)** PCA plot showing the clustering of C10 and SVEC cells based on scProteomics data. Total 1,141 proteins were used in the PCA projection. **(D)** Box plot showing the distributions of gene identification numbers for QC-passed 65 C10 cells and 74 SVEC cells. Centerlines represent the distribution median and “black” points represent the distribution mean. **(E)** Violin plots showing the coefficient of variations of protein abundances from QC-passed scRNAseq samples processed using nanoSPINS (n = 8,423 genes for C10 and n = 8,118 genes for SVEC cells). “Black” point represents median CV. **(F)** PCA plot showing the clustering of C10 and SVEC cells based on scRNAseq data. Total 12,722 genes were used in the PCA.

Regarding scRNAseq, we evaluated the transferred single-cell droplets, comprising 96 C10 and SVEC cells each, as well as 24 control and blank samples. To address the systematic and random noise in scRNA-seq datasets that can interfere with biological interpretation, we applied several QC metrics ^34^. By filtering with a minimum of 50% mapped reads and at least 1,500 genes/features per cell, we retained a total of 139 high-quality cells, consisting of 65 C10 and 74 SVEC cells. Similarly, the threshold of 1500 genes per cell was established based on the number of genes detected in the control samples (**Supplementary Figure 5A**). On average, unique molecular identifier (UMI) counts of approximately 12,411 and 8,905 were observed from high quality C10 and SVEC cells, respectively (**Supplementary File 3**). Additionally, mean number of detected genes per cell was approximately 3,796 for C10 and 3,515 for SVEC cells (**Figure 4D**). Using a raw read cut-off of 1, and a minimum of 5 cells per cell type observing a feature, a total of 12,722 unique genes were identified across all single-cells. Of these, 9,006 genes were common to both cell-types, with 1,929 genes unique to C10 and 1,787 unique to SVEC (**Supplementary Figure 5B**). Additionally, the median percentage of mitochondrial reads captured from C10 and SVEC cells using the nanoSPINS workflow was 3.57% and 2.16%, respectively (**Supplementary Figure 5C**), indicating high cell quality and minimal cellular stress (e.g., apoptosis or cell death), which supports the reliability of downstream biological interpretations. To assess the impact of the nanoSPINS workflow on the transcriptome profiles, we compared the nanoSPINS cells to 72 directly-sorted control C10 cells and 72 directly-sorted SVEC cells. We observed that the average number of genes identified per cell was nearly two-fold higher in intact cells, with 7,463 and 6,478 genes identified from 64 high quality C10 cells and 68 high quality SVEC cells, respectively (**Supplementary Figure 5D** and **Supplementary File 4**). This is not unanticipated, as nanoSPINS cells retain intact nuclei on the chip after transferring, reducing the amount of transferred mRNA to the cellenCHIP. Across all high-quality directly sorted cells, a total of 15,589 unique genes were identified. Of these, 12,545 genes were shared between directly sorted cells and those processed with the nanoSPINS workflow, while 3044 genes were unique to directly-sorted cells, and only 177 genes were unique to nanoSPINS-processed cells (**Supplementary Figure 5E**). Overall, 98.61% of genes detected using nanoSPINS workflow were also represented in the transcriptomic profile captured from directly sorted cells, further demonstrating that the nanoSPINS workflow does not introduce significant transcriptomic artifacts.

Median CV values for single-cell transcriptomic measurements captured via the nanoSPINS workflow were 0.41 in C10 cells and 0.35 in SVEC cells (**Figure 4E**). The increased variation (relative to the scProteomics) reflects characteristics of mRNA expression, such as lower copy numbers and shorter half-lives. The median CV values for directly sorted C10 and SVEC cells were both 0.42, indicating the nanoSPINS workflow did not introduce additional variation into the scRNAseq datasets (**Supplementary Figure 5F**). Furthermore, PCA separation of cell types could be achieved along PC2 and PC4 for the nanoSPINS-transferred cell lysates, with both components showing significant (pvalue < 0.001) correlation with cell type, respectively (**Figure 4F**). Pearson correlation analysis revealed average correlation coefficient (r) values of 0.65 and 0.64 for the scRNA-seq datasets generated from C10 and SVEC cell-types, respectively, using nanoSPINS platform (**Supplementary Figure 6**). Similarly, average r values of 0.62 and 0.65 were observed for scRNAseq datasets generated from directly sorted C10 and SVEC cell-types, respectively. These correlation values are consistent with previously reported ranges for scRNA-seq datasets^14, 35^, and suggest that the quality of the transcriptome captured by the nanoSPINS platform is on par with that of the directly sorted cells.

Comparison of the two modalities revealed that 89.8 % of the proteins identified from high-quality cells were represented in the transcriptome captured using the nanoSPINS platform (**Figure 5A**). Differential abundance analysis of the transcriptomic and proteomic profiles between C10 and SVEC cells revealed features enriched for each cell type (**Supplementary File 5**). Interestingly, the overlap of significant features at mRNA (n = 165; abs(log2FC) >= 1, padj < 0.05) and protein (n = 64; abs(log2FC) >= 0.5, padj < 0.05) levels was relatively low, with only 10 transcript-protein pairs shared showing similar expression/abundance profiles (**Figure 5B**). This suggests that, even if statistically significant, most cell-type markers identified using a single modality may not be reliably observed across other modalities. Previous studies on C10 and SVEC cells have established the expression profiles of various genes at both protein and transcript levels, such as H2-K1 and H2-D1 in SVEC cells and COL1a1, COL3a1 and FN1 in C10 cells^14, 36^. Our findings corroborate these studies and help to validate the current approach, showing that these genes are significantly more abundant in SVEC and C10 cells, respectively, at both the protein and mRNA levels (**Figure 5C**). Additionally, we found that several previously identified cell type-specific surface marker proteins were significantly (padj < 0.05) more abundant in SVEC and C10 cells, respectively (**Supplementary File 5**). For example, HSP90aa1 and BST2 were enriched in SVEC cells, whereas PDIA6, EZR, and F11R (also known as JAM1) showed higher abundance in C10 cells^18^. Taken together, these results provide strong evidence that the nanoSPINS-based single-cell multiomics platform can provide sensitive and reproducible quantitative measurements of both the transcriptome and proteome of the same single cells.

**Figure 5.**
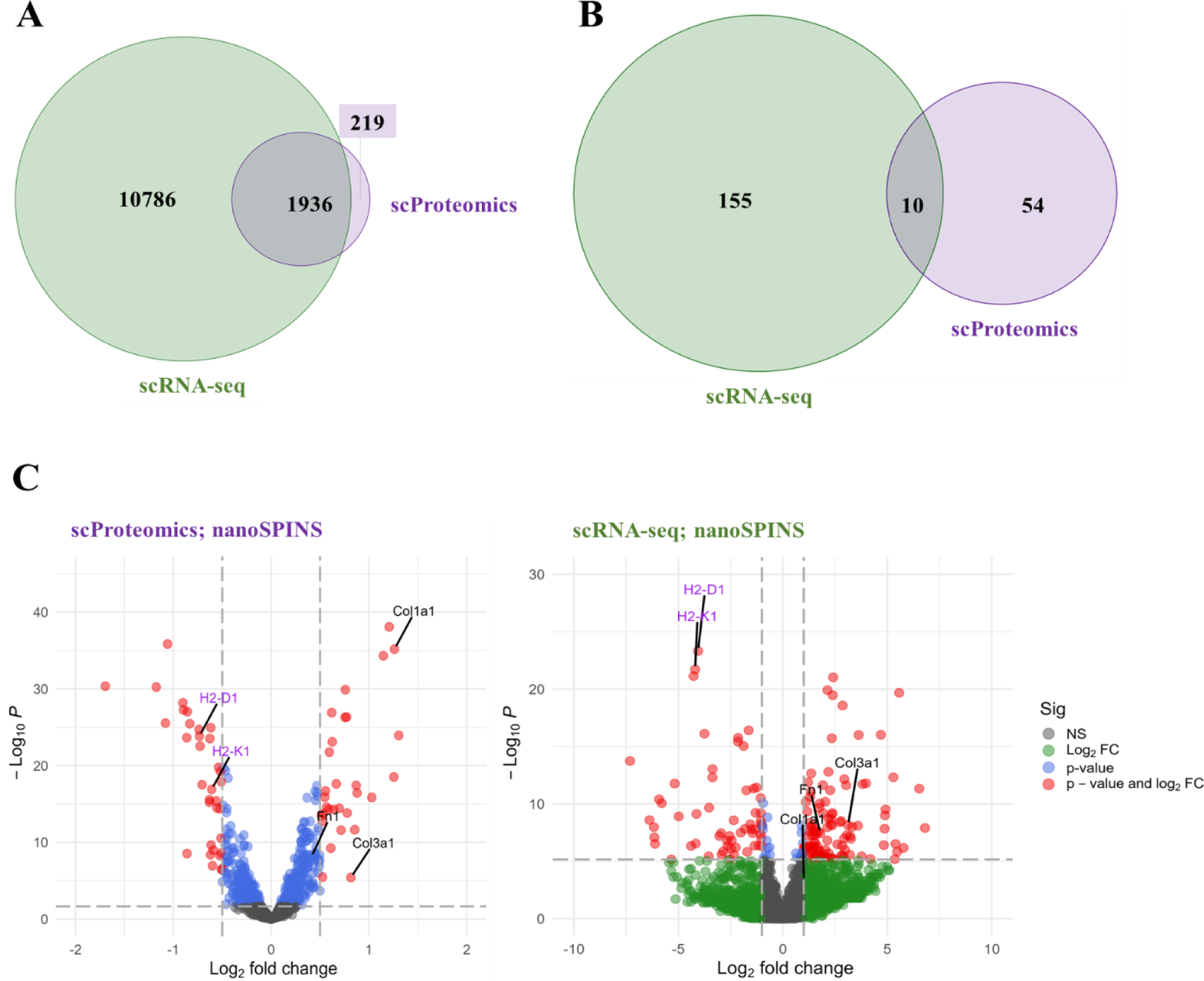
**(A)** Venn diagram showing the overlap between high-confidence genes and proteins identified from quality controlled single cells. Dark green represents genes from scRNA-seq analysis, and Indigo represents proteins from single-cell proteomics. **(B)** Venn diagram illustrating the overlap between significantly differentially expressed genes (|log₂FC| ≥ 1, adjusted p < 0.05) and proteins (|log₂FC| ≥ 0.5, adjusted p < 0.05) from the comparison between C10 and SVEC single cells. Dark green and indigo represents significant differentially expressed genes and proteins in C10 cells vs SVEC single cells identified from scRNA-seq and scProteomics analysis. **(C)** Volcano plots showing the distribution of significantly differentially expressed genes and proteins between C10 and SVEC cells. Significance thresholds were set at |log₂FC| ≥ 1 (for genes) and |log₂FC| ≥ 0.5 (for proteins) with adjusted p < 0.05. P-values were corrected for multiple testing using the Benjamini-Hochberg method.

## DISCUSSION

To enable high-throughput, global transcriptomic and proteomic measurements at the single-cell level, we first sought to better understand the mechanisms of cell lysis in nanoSPLITS. We determined that the presence of a detergent such as DDM can have a significant impact on cell lysis in a hypotonic buffer. In the absence of DDM, osmotic stress plays a larger role in cell lysis, while the inclusion of DDM permeabilizes the cell membrane, allowing for the transfer of diffusible cytosolic molecules into the nanodroplet. Interestingly, in both cases, cell lysis remains incomplete and largely unaffected by a single freeze-thaw. Considering this observation and recognizing that integration of the cellenCHIP 384^31^ would require a new method of transferring cellular components, we wondered if centrifugation might facilitate such an effect. Towards this end, we designed a nested nanoPOTS chip to match the cellenCHIP 384 well arrangement and demonstrated how evaporation of the lysis buffer prior to reconstitution with water could ensure the retention of cellular material for TMTpro-based scProteomics while allowing for centrifugation-enabled transfer of soluble components to the cellenCHIP 384 for scRNAseq.

This nanoSPINS approach increased the throughput for performing multiomic measurements from same single cells by 8-fold as compared to our previous nanoSPLITS platform^14^. With the introduction of higher multiplexing capabilities, such as TMT-32plex^16^, we anticipate that the throughput of this workflow can be improved even more.

From a set of 96 C10 and 96 SVEC cells analyzed through a single nanoSPINS transfer, approximately 1,186 proteins and 3,646 genes were detected per single cell. Average Pearson correlations of ∼0.94 and ∼0.64 across single cells for scProteomics and scRNAseq, respectively, demonstrate that the nanoSPINS workflow can deliver high-quality data comparable to published benchmarks^14, 18, 35–37^. Moreover, median CVs at the protein and gene level were 0.27 and 0.38, respectively, across both cell types, further underscoring the workflow’s reproducibility. These results were comparable to prior scProteomics and scRNAseq workflows, despite sample handling being a known source of technical variability^38, 39^. Prior studies have also applied scProteomics and scRNAseq to C10 and SVEC cells, providing a form of validation for cell-enriched proteins for each cell-type. Not only were we able to identify the majority of the cell-type enriched proteins for C10^18, 36^ (such as COL1A1, EZR) and SVEC^14, 36^ (such as H2-K1, BST2) cells using nanoSPINS workflow, but also demonstrated their cell-specific enrichment at the transcript level. Similar to previous reports, we also observed limited overlap between cell-type enriched proteins and genes, likely due to the low correlation between the two modalities, suggesting that mRNA expression does not necessarily reflect protein abundance in this context. However, certain overlapping protein–gene pairs exhibited consistent enrichment trends, such as H2-K1 and H2-D1 in SVEC cells, and Col3a1 in C10 cells, at both the protein and transcript levels. These findings underscore the importance of parallel multimodal measurements for high-confidence biomarker discovery and cross-modality applications, helping to validate the nanoSPINS approach and underscore the promise of multimodal single-cell profiling in the context of biomarker discovery.

## Supporting information

Supplementary_Word_Document

## DATA AVAILABILITY

The mass spectrometry raw data have been deposited to the ProteomeXchange Consortium via the MassIVE partner repository with dataset identifiers MSV000099357. The raw RNAseq data is available at NCBI under accession PRJNA1337717. Scripts, result files, and associated metadata used are available at the nanoSPINS_data_scripts GitHub repository [https://github.com/prdawar/nanoSPINS]

## CODE AVAILABILITY

Associated code for the data analysis is included in R markdown or R script files at the GitHub repository [https://github.com/prdawar/nanoSPINS]

## AUTHOR CONTRIBUTIONS

P.D.: Formal Analysis; Visualization; Writing - Original Draft, Reviewing and Editing. L.M.M: Methodology; Investigation; Writing - Reviewing and Editing. S.M.W: Investigation. H.D.M: Data Curation. J.W.B: Methodology; Data Curation. J.C.B.: Conceptualization; Methodology; Data Curation; Funding acquisition; Writing - Reviewing and Editing. A.S.: Methodology; Writing - Reviewing and Editing. C.C.B: Resources; Methodology. L. P-T.: Supervision; Project administration; Funding acquisition; Writing - Reviewing and Editing. Y.Z.: Supervision; Conceptualization; Investigation; Methodology; Project administration; Writing - Reviewing and Editing. J.M.F.: Conceptualization; Investigation; Methodology; Project administration; Formal Analysis; Visualization, Writing - Original Draft, Reviewing, and Editing. The final manuscript was read and approved by all authors. The final manuscript was read and approved by all authors.

## ACKNOWLEDGMENTS

We thank Dr. Matthew Monroe for his assistance in depositing the raw proteomic data into MassIVE. This work was supported by an Environmental Molecular Sciences Laboratory project award (10.46936/prtn.proj.2021.51834/60000340, J.C.B.) and a Pacific Northwest National Laboratory Cooperative Research and Development Agreement (CRADA) with Scienion (10.46936/staf.proj.2023.60714/60008812, J.M.F. and L.P-T.). This research was performed at the Environmental Molecular Sciences Laboratory, a DOE Office of Science User Facility sponsored by the Biological and Environmental Research program under Contract No. DE-AC05-76RL01830.

