## Supplementary_Word_Document for "High-throughput Single-Cell Proteomics and Transcriptomics from the Same Cells with a Nanoliter-Scale Spin-Transfer Approach"

^3^Present address: Department of Proteomic and Genomic Technologies, Genentech, 1 DNA Way, South San Francisco, 94080, United States

^*^Correspondence

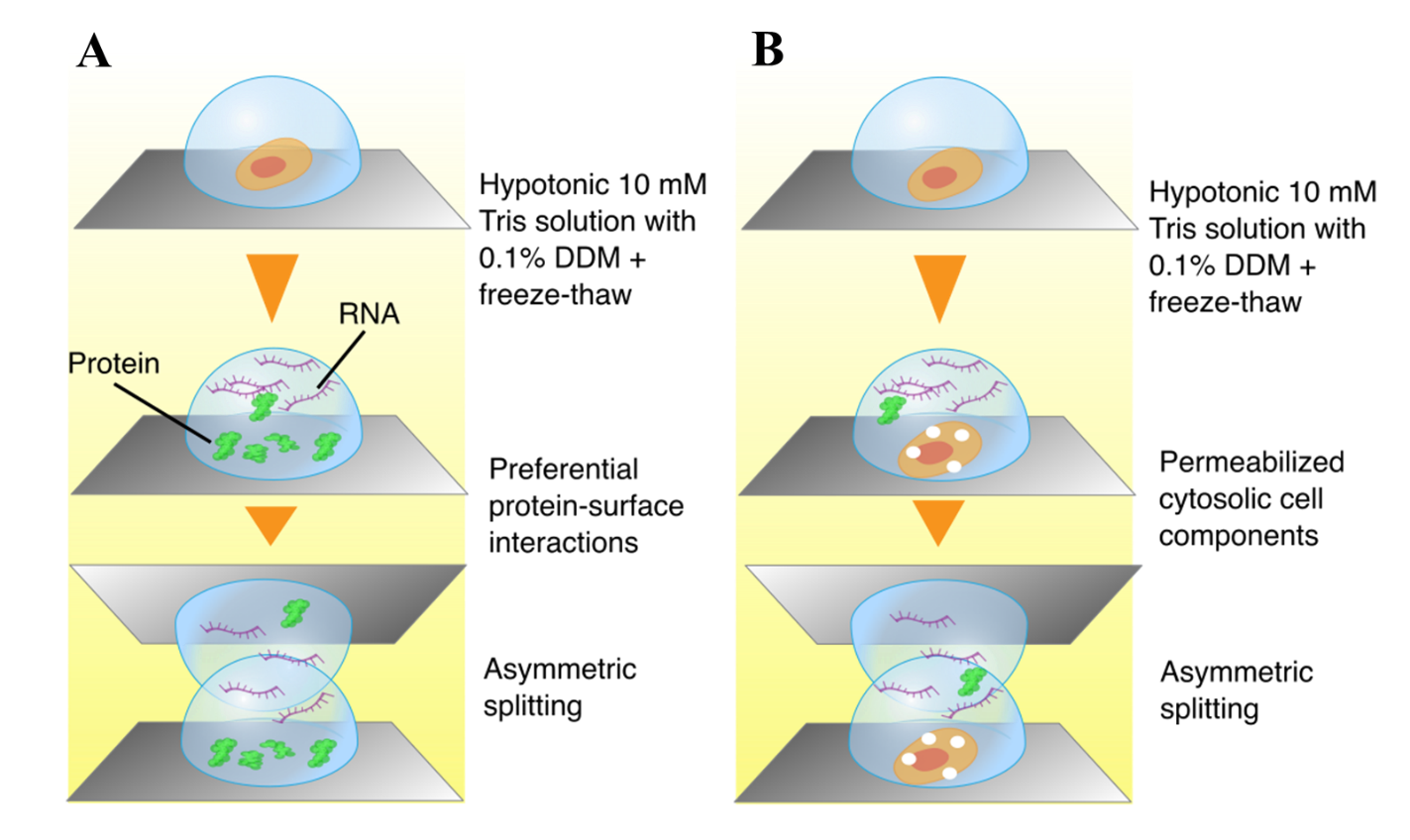

**Supplementary Figure 1** – **(A)** Previous model of cell lysis - 0.1% DDM-mediated increased overall cell lysis, with enhanced protein retention due to hydrophobic interactions between protein molecules and the trimethylsilyl-coated surface of the nanoSPLITS chip wells. **(B)** Revised model suggesting that 0.1% DDM primarily induces membrane permeabilization rather than complete lysis, allowing diffusible cytosolic components to enter the droplet while the bulk of the cell remains intact and adhered to the glass chip surface.

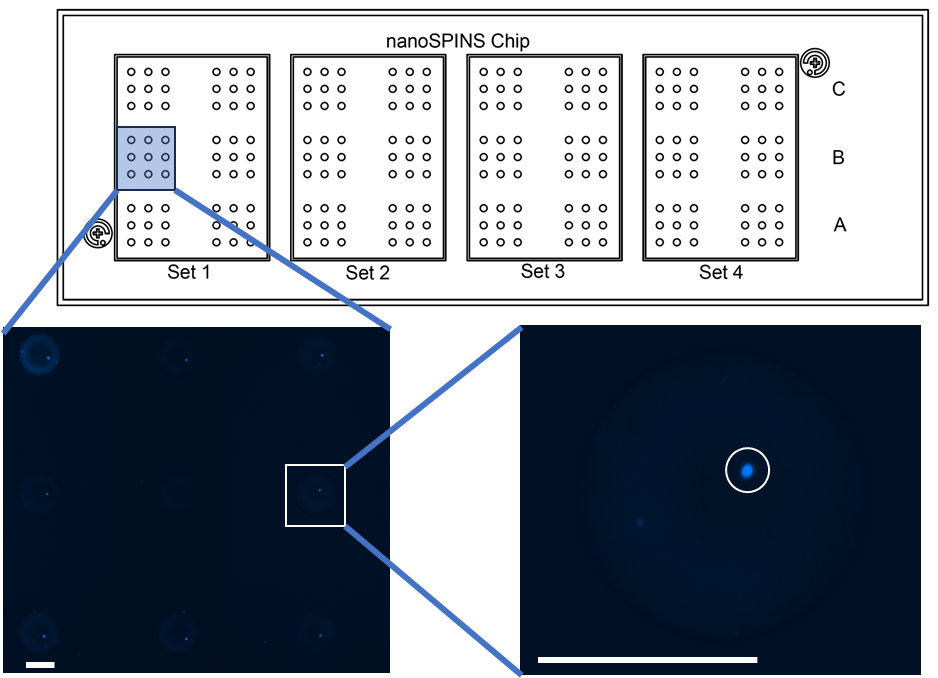

**Supplementary Figure 2** – Fluorescent image of a 3×3 array of wells on the nanoSPINS chip, showing eight peripheral wells containing Hoechst-stained single cells sorted into 40 nL of 0.1% DDM, 10 mM HEPES (pH 8.5), and a central control/empty well, demonstrating high cell retention on the nanoSPINS wells following a single freeze-thaw cycle, evaporation, immediate reconstitution with 40 nL nuclease-free water, and centrifugation at 1000 × g.

**Set 1**

**Set 2**

**Set 3**

**Set 4**

**Supplementary Figure 3** – Arrangement of samples on cellenCHIP 384 used for scRNAseq sample preparation. Green, Blue and Grey wells represents C10, SVEC cells and Blanks (negative control) from nanoSPINS chip, respectively. Orange and Red wells represents C10 and SVEC cells directly sorted onto cellenCHIP, respectively. White wells represents empty wells (negative control) for directly sorted cells.

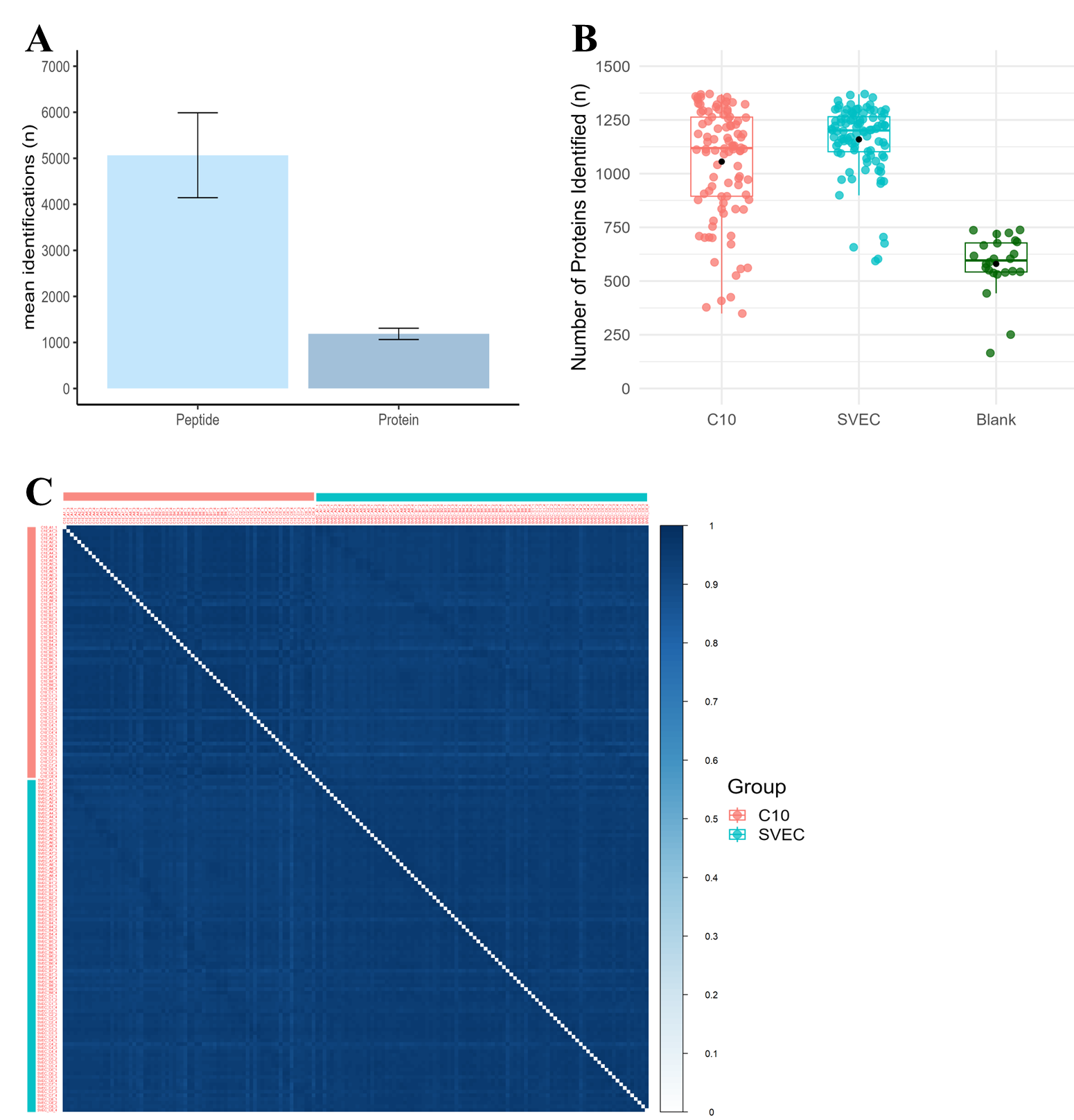

**Supplementary Figure 4** – **(A)** Mean number of detected peptides and proteins from a total 160 QC-passed single cells (69 C10 and 91 SVEC). Error bars indicate standard deviations ( ± s.d.). **(B)** Box plot showing the distributions of protein identification numbers for 96 C10 cells, 96 SVEC cells and 24 blanks. Centerlines represents the medians and “black” points represents the mean. **(C)** Clustering matrix showing Pearson correlations across 160 single cells using log2-transformed protein intensities. The color scale indicates the range of Pearson correlation coefficients.

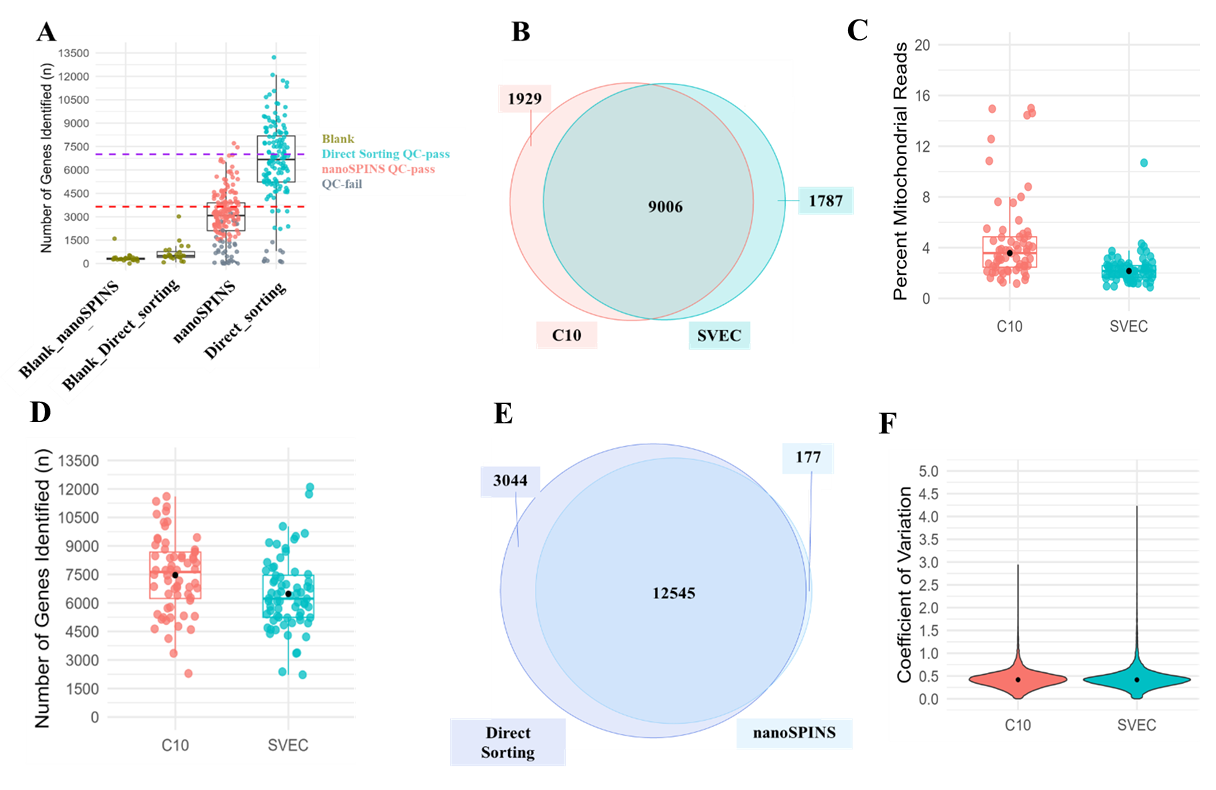

**Supplementary Figure 5** – **(A)** Box plot showing the distributions of numbers of genes identified from 96 C10 cells, 96 SVEC cells and 24 blank/control processed using nanoSPINS platform and 72 SVEC cells and 72 C10 cells sorted directly and respective 24 blank/control. “QC_fail” cells represents cells with less than 1500 gene ids and less than 50% of the mapped reads overall. Red and Purple dash line represents mean number of genes identified in high quality nanoSPINS cells and directly sorted cells. **(B)** Venn diagram showing the overlap between high-confidence genes identified from quality controlled C10 (presented in red) and SVEC (presented in Strong cyan) cells processed using nanoSPINS workflow. **(C)** Box plot showing the distributions of number of reads mapped to mitochondrial genes in QC-passed C10 cells and SVEC cells. Centerlines and “black” points represents the distribution median. **(D)** Box plot showing the distributions of gene identification numbers for QC-passed 68 C10 cells and 64 SVEC cells sorted directly. Centerlines represents the distribution median and “black” points represents the distribution mean.  **(E)** Venn diagram showing the overlap between high-confidence genes identified from quality-controlled cells processed using nanoSPINS workflow (n = 139; LightSkyBlue) and cells sorted directly (n = 132; RoyalBlue). **(F)** Violin plots showing the coefficient of variations of gene expression from QC-passed scRNAseq samples sorted directly (n = 14,759 genes for C10 and n = 14,482 genes for SVEC cells). “Black” point represents median CV.

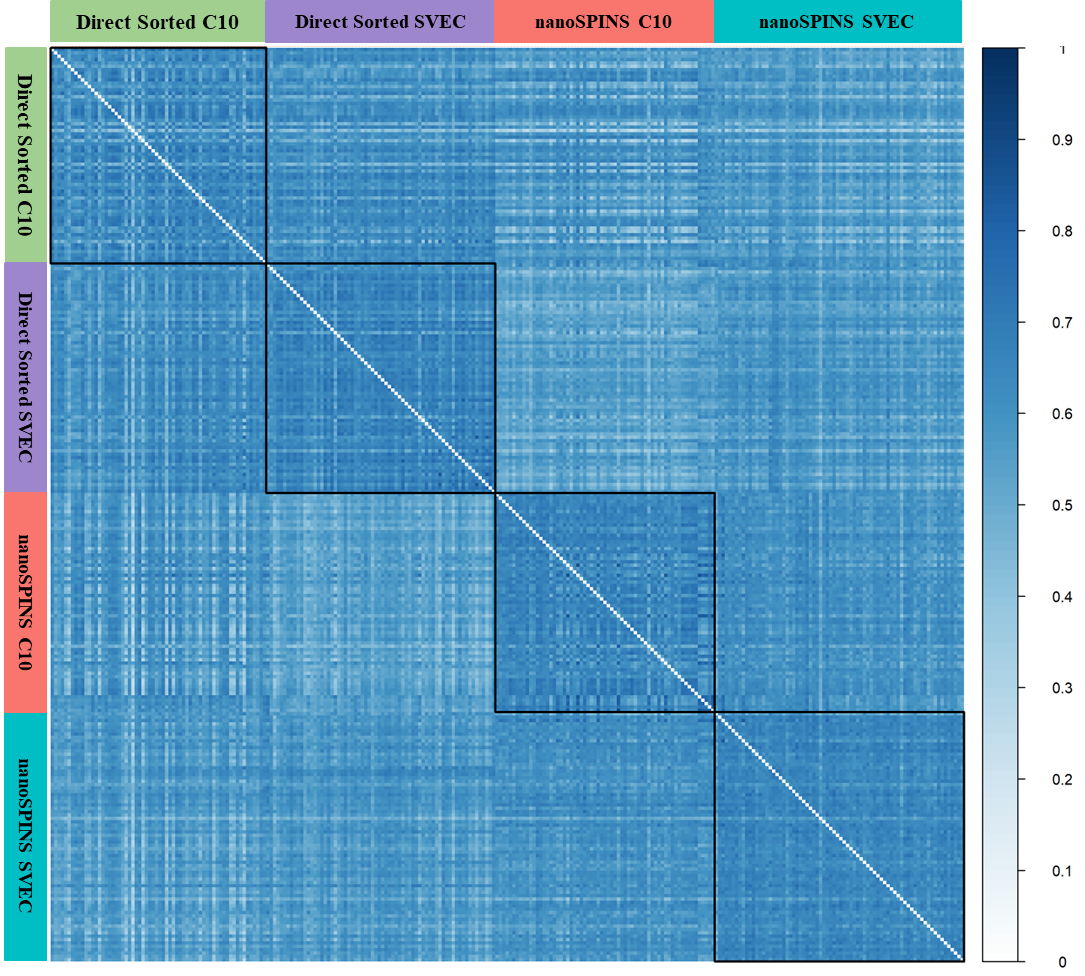

**Supplementary Figure 6** - Clustering matrix displays pairwise Pearson correlation coefficients among 271 QC-passed single cells, including 74 SVEC and 65 C10 cells processed via the nanoSPINS workflow, and 68 SVEC and 64 C10 cells sorted directly onto the nanoSPINS chip. The color scale indicates the range of Pearson correlation coefficients.
